# PanGraph: scalable bacterial pan-genome graph construction

**DOI:** 10.1101/2022.02.24.481757

**Authors:** Nicholas Noll, Marco Molari, Liam P. Shaw, Richard A. Neher

## Abstract

The genomic diversity of microbes is commonly parameterized as single nucleotide polymorphisms relative to a reference genome of a well-characterized, but arbitrary, isolate. However, any reference genome contains only a fraction of the microbial *pangenome*, the *total* set of genes observed in a given species. Reference-based approaches are thus blind to the dynamics of the accessory genome, as well as variation within gene order and copy number. With the wide-spread usage of long-read sequencing, the number of high-quality, complete genome assemblies has increased dramatically. Traditional computational approaches towards whole-genome analysis either scale poorly with the number of genomes, or treat genomes as dissociated “bags of genes”, and thus are not suited for this new era. Here, we present *PanGraph*, a Julia-based library and command line interface for aligning whole genomes into a graph. Each genome is represented as an undirected path along vertices, which in turn, encapsulate homologous multiple sequence alignments. The resultant data structure succinctly summarizes population-level nucleotide and structural polymorphisms and can be exported into a several common formats for either downstream analysis or immediate visualization.

---

During evolution, microbial genomes change by both local mutations and large-scale alterations (Arnold *et al*., 2021). Local mutations only change a few nucleotides by substitution, insertion or deletion. Conversely, large-scale alterations reorganize the sequence, and involve either the homologous recombination of large segments, gene loss or gain, inversions, or mobilisation of genetic elements. The accumulation of such changes over time complicates comparative genomic analyses of present day isolates. Homologous recombination is rapid enough that most genes in many bacterial core genomes have distinct phylogenies (Sakoparnig *et al*., 2021) and even closely related genomes differ dramatically in gene content.

Recent advances in long-read sequencing have enabled the low-cost assembly of complete genomes at the quality of reference databases (Whibley *et al*., 2021). The accumulation of so many complete genomes promises to rapidly improve our ability to quantify the evolutionary dynamics that drive microbial diversity in natural populations. Since reference-based approaches only partially capture microbial diversity (Tettelin *et al*., 2008), the concept of the *pangenome* has motivated the development of methods that account for substantial variation in gene content. However, though such approaches can accurately capture nucleotide polymorphisms within genes, they approximate structural polymorphisms as gene presence-absence relationships (Ding *et al*., 2018; Page *et al*., 2015) irrespective of gene order or orientation. This new era of pangenomics demands novel data structures to encapsulate the *complete* diversity of a given genomic sample set.

In recent years, efforts have focused on generalizing the *pangenome* framework of microbial diversity to graphical models (Eizenga *et al*., 2020). At a high level, pangenome graphs generalize the reference sequence coordinate system conventionally used and encode genomes as directed paths through the graph consisting of node that represent sequence. In multicellular eukaryotes, pangenome graphs usually have an overall linear structure in which structural diversity can be encoded as short range excursions. In microbial genomics, in contrast, large scale rearrangements and inversions give rise to complex graph topologies. Here, we focus on pangenome graphs of closely related bacterial genomes.

While easy to conceptualize, the construction of pangenome graphs has proven computationally challenging. Colored generalizations of the de Bruijn graph-based assemblers have been successively used to build graphs from large sequence sets, although the underlying efficiency derives from a fixed k-mer size which prevents modelling long-range homology (Iqbal *et al*., 2012; Muggli *et al*., 2017). An orthogonal approach has been to formulate the inference of the pangenome graph as a multiple genome alignment. However, current methods either scale poorly to large sets of genomes (Darling *et al*., 2010), focus on comparisons across diverse sets of species across the tree of life at the cost of memory (Armstrong *et al*., 2020), or utilize a reference-guided approach by partitioning genomes first into annotated genes (Colquhoun *et al*., 2021; Gautreau *et al*., 2020).

Here we present *PanGraph*, a Julia (Bezanson *et al*., 2017) library and command line interface, designed to efficiently align large sets of closely related genomes into a pangenome graph on personal computers. The resulting graph both compresses the input sequence set and succinctly captures the population diversity at multiple scales: from nucleotide mutations and indels to structural polymorphisms driven by inversions, rearrangements, and gene gain/loss. The underlying graph data structure can be exported into numerous formats for downstream analysis and visualization in software such as Bandage (Wick *et al*., 2015).

## I. ALGORITHMS AND IMPLEMENTATION

*PanGraph* transforms an arbitrary set of genomes into a *graph* that simultaneously compresses the collection of sequences and exhaustively summarizes both the structural and nucleotide-level polymorphisms. The graph is composed of *pancontigs*, which represent linear multiple-sequence alignments of homologous sequence found within one or more input genomes. *Pancontigs* are connected by an edge if they are syntenic on at least one input sequence; individual sequences are then recapitulated by contiguous *paths* through the graph. Pancontigs are directed, meaning their orientation in a path can be either 5’ to 3’ or vice versa.

To construct a pangraph, the algorithm needs to find homologous sequence within and among all input genomes. *PanGraph’s* overarching strategy is to approximate multiple-genome alignment by iterative pairwise alignment of graphs of subsets of sequences, in the spirit of progressive alignment tools (Armstrong *et al*., 2020; Darling *et al*., 2010; Feng and Doolittle, 1987). Pairwise graph alignment is performed by an all-to-all alignment of the *pancontigs* between both graphs and the order of pairwise alignments is determined by a guide tree.

### A. Guide tree construction

The alignment guide tree is constructed subject to three design constraints: (i) similar sequences are aligned first, (ii) the similarity computation scales subquadratically with the number of input sequences, and (iii) the resultant tree is balanced to maximize parallelism. To this end, we formulate the algorithm as a two step process. The initial guide tree is constructed by neighbor-joining (Saitou and Nei, 1987); the pairwise distance between sequences is approximated by the Jaccard distance between sequence minimizers (Roberts *et al*., 2004). Computationally, each sequence can be sketched into its set of minimizers in linear time while the cardinality of all pairwise intersections can be computed by sorting the list of all minimizers to efficiently count overlaps. Hence, the pairwise distance matrix is estimated in a time that increases log-linearly in the total length of sequence that is analyzed. The final guide tree is constructed as the balanced binary tree constrained to reproduce the topological ordering of leaves found initially. This balancing maximizes the number of independent pairwise alignments and thereby allows efficient parallelism. See Fig. 1A for a graphical depiction of the guide tree.

**FIG. 1.**
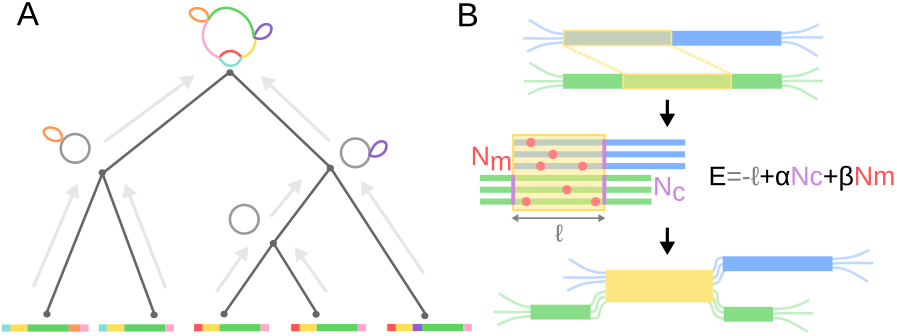
Overview of PanGraph algorithm. (A) The alignment graph is constructed progressively by aligning graphs pairwise up a guide tree constructed from neighbor-joining the minimizer overlap between strains. (B) During pairwise alignment, *pancontigs* (blue and green) are merged by identifying homologous intervals (shown in yellow). If the underlying alignments are viewed compatible, i.e. the energy is less than 0, the pancontigs are merged.

### B. Iterative graph alignment

The full pangraph representing all genomes is constructed by aligning/merging graphs that represent subsets of the genomes in an iterative manner illustrated in Fig. 1. The iteration starts with one subgraph per input genome, each representing its respective genome as a single *pancontig*. Pairs of subgraphs are aligned in a postorder traversal of the guide tree. The identified homologous intervals of the pancontig are then merged, thereby creating shorter contigs that represent homologous sections of multiple input genomes, see Fig. 1B for a graphical depiction. These steps of pairwise graph alignment and merging of homologous intervals are repeated until the root of the guide tree is reached. *Pancontigs* encapsulate linear multiple-sequence alignments which are modelled internally by a star phylogeny, i.e. are assumed to be well-described by a reference sequence augmented by independent SNPs and indels for each contained isolate.

#### Pairwise alignment

To align two graphs, the consensus sequences of all pancontigs in both graphs are searched for homologies and aligned. Full genome alignment between two closely related isolates is a well-studied problem with many sensitive and efficient tools available (Li, 2018; Marçais *et al*., 2018). We chose to use *minimap2* as the core pairwise genome aligner for its proven speed, sensitivity, and easy-to-use exposed library API (Li, 2018). This alignment kernel is included within a custom Julia wrapper, available at github.com/nnoll/minimap2_jll.jl. However, we note that *PanGraph* is written to be modular, and additional alignment kernels can be added with ease. In particular we decided to include the option to use *mmseqs2* (Steinegger and Söding, 2017) as an alternative alignment kernel, because of its sensitivity on highly diverged sequences at the cost of higher computational time.

#### Merging of homologous sequence

If the above alignment step detected homologous stretches between two pancontigs, these pancontigs, or parts of them, can be merged. It is not uncommon that one pancontig has homology with multiple other pancontigs and the iterative algorithm has to make choice which potential mergers are performed and in which order. We rank each alignment between two pancontigs according to the pseudo-energy

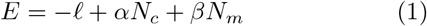

where *ℓ, N*_*c*_, and *N*_*m*_ denote the alignment length, number of *pancontigs* created by the merger, and number of polymorphisms per genome in the newly created pancontig respectively. See Fig 1B for a graphical definition for each term. Additionally, *α* and *β* are hyperparameters of the algorithm. Only mergers whose alignment has negative pseudo-energy are performed. The parameter *β* controls the maximal sequence divergence between the two merger-candidates and no mergers will be performed if the Hamming distance exceeds 1*/β*. Importantly, this divergence threshold is applied to the comparison of the consensus sequences of the merger candidates. The parameter *α* controls the fragmentation of the graph and imposes a minimal length on the resulting pan-contigs resulting from the merger. The merger is always performed if the merger doesn’t increase the number of contigs, but mergers that introduce 1, 2, 3 or 4 cuts are only performed for sufficiently long homologous stretches as parameterized by *α*. In addition, there is parameter that controls the minimal size of contigs produced by the algorithm.

At the graph level, the merger of two *pancontigs* defines a new *pancontig*, connected on both sides by edges to the neighboring *pancontigs* of both inputs, and thus locally collapses the two graphs under consideration. At the nucleotide level, the pairwise alignment of two *pancontigs* maps the reference of one onto the other; the merger of two *pancontigs* requires the application of the map onto the underlying multiple-sequence alignment. Once both sets of sequences are placed onto a common coordinate system, the resultant consensus sequence, and thus polymorphisms, are recomputed. This procedure can be viewed as an approximate multiple sequence alignment algorithm that aligns homologous parts without further investigating insertions in the two sub-alignments.

The above procedure is repeated until no alignments with negative energy remain. Upon completion, transitive edges within the graph, i.e., edges along which all path are co-linear, are removed by merging adjacent *pancontigs*.

### C. Parallelism

*PanGraph* guide trees are balanced binary trees that break the task into many independent sub-problems and thereby enable scalable parallelism, as shown in Fig 1A for a cartoon example. Each internal node of the guide tree represents a job that performs a single pairwise graph alignment between its two children. The process will block until both of its children processes have completed and subsequently pass the result of their pairwise graph alignment up to the parent. All jobs run concurrently from the start of the algorithm; the Julia scheduler resolves the order of dependencies naturally (Bezanson *et al*., 2017). As such, the number of parallel computations is automatically scaled to the number of available threads allocated by the user at the onset of the alignment.

### D. Graph Export and Availability

The constructed pangenome graph can be exported in a variety of file formats for downstream analysis and visualization. In addition to a custom JSON schema, PanGraph can export the alignment as a GFA file, where each *pancontig* is represented as a segment and each genome as a path. This allows for visualization in software such as Bandage (Wick *et al*., 2015). Lastly, we provide functionality to export as a conventional presence/absence pangenome – albeit one with *pancontigs* taking the place of putative gene clusters – that can be visualized directly by the PanX toolkit (Ding *et al*., 2018).

PanGraph is published under an MIT license with source code, extensive documentation, examples, and instructions for installation available at github.com/neherlab/pangraph. Additionally, pre-built binaries are available as GitHub releases. All data and scripts used to validate PanGraph are available within the same repository.^1^

## II. VALIDATION AND PERFORMANCE

### A. Validation on synthetic data

As a first validation step we use generated synthetic data to quantify the performance characteristics of *Pan-Graph* as a function of input size, and its accuracy as a function of sequence diversity.

#### Generation of synthetic data

We simulated populations of size *N* = 100 and varying sequence length up to *L* = 500 000 utilizing a Wright-Fisher model (Hudson, 2002) evolved for *T* = 50 generations. In addition to nucleotide mutations that occur at rate *μ* per generation, we modelled inversions, deletions (respective rates 0.01 and 0.05 per generation), and horizontal sequence transfer that occurs with tunable rate *h* per generation per genome (see SI Section I for details). The ancestral state for each sequence is tracked through each evolutionary event so that the true mosaic relatedness structure can easily be converted to a graph. The simulation framework is distributed within the PanGraph command line tools for external use.

#### Performances as a function of dataset size

The algorithmic complexity was measured empirically by constructing pangenome graphs from data generated by simulating populations of increasing size and average sequence length. The mean and standard deviation of the run-time obtained from 50 iterations are shown in Fig. 2. Importantly, PanGraph’s computational complexity grows *linearly* with the number of input genomes once the number of genomes exceeds a certain threshold and parallelism can be exploited. We note that PanGraph scales approximately log-linear with average sequence length, as expected from the underlying algorithmic complexity of *minimap2* (Li, 2018). Benchmarks of CPU and memory usage are reported in SI Section IIA.

**FIG. 2.**
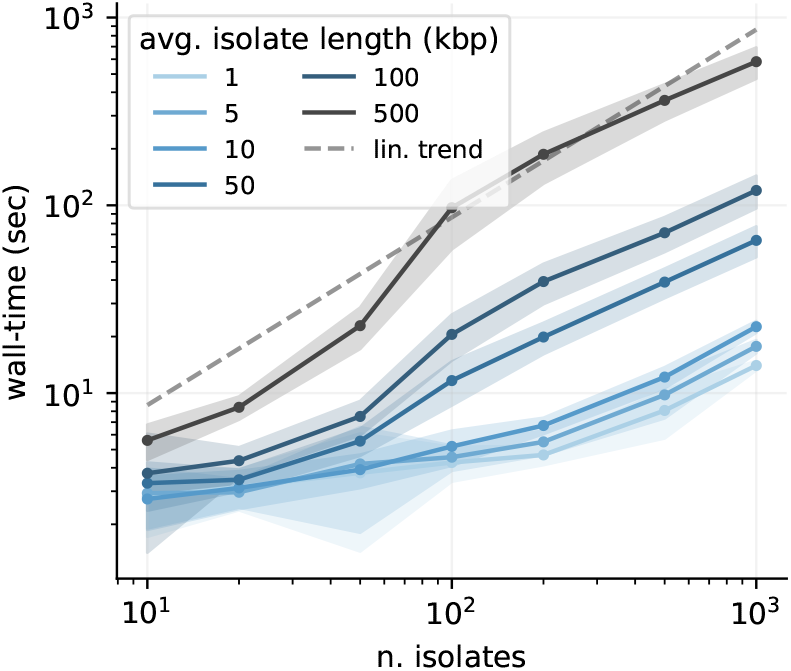
Algorithm performance. PanGraph scales linearly with the number of input genomes. This is a direct result of the guide tree simplification. The solid line and ribbons display the mean and standard deviation over 50 runs. All runs were performed utilizing 8 cores, and with the default *minimap2* alignment kernel and *asm20* option.

#### Accuracy as a function of sequence divergence

To be useful and informative beyond data compression of microbial genomes, the contigs and break-points of the reconstructed graphs should correspond, at least approximately, to evolutionary events in the history of the sample. We quantified PanGraph’s ability to accurately reconstruct the true graph by computing the displacement of the inferred breakpoints relative to their known locus stored from the evolutionary simulation (cf. SI Section IIB for details). We evaluate this displacement by generating datasets with different rates of mutation *μ* and horizontal sequence transfer *h*. For each (*h, μ*) pair we perform 25 different simulations, and build pangenome graphs using three different options for the alignment kernel: *minimap2* (Li, 2018) *with asm10* and *asm20* options and *mmseqs2* (Steinegger and Söding, 2017). *Critically, we found that the accuracy was independent* of the rate of horizontal transfer *h* and thus underlying graph complexity. The predominant determinant of accuracy was the sample diversity, controlled by the mutation rate *μ* in our simulations (cf. SI Fig. 2). For low-diversity datasets, breakpoints are inferred with accuracy of few basepairs, while for highly diverged isolates most breakpoints are displaced by several hundreds of basepairs (cf. SI Fig. 3). The choice of alignment kernel influences the threshold diversity at which accuracy is lost, suggesting that inability to detect homology between diverged sequences is the reason.

For each alignment kernel and value of average sequence divergence in the population we evaluated the fraction of breakpoints that have displacement greater than the default minimal block size for Pangraph of 100 bp (cf. Fig. 3). On the sequences generated by our simulations, *minimap2* with option *asm20* show a loss of accuracy at an average pairwise sequence divergence of around 5%, while *mmseqs2* is accurate up to around 12%, at the cost of higher computational time. These threshold are lower than the nominal sensitivity of the aligners, which are 10% and 20% for minimap2 with settings *asm10* and *asm20*, respectively, and around 30% for *mmseqs2*. This discrepancy is due to the fact that for accurate graph constructions, the largest divergence, rather than the average divergence, is the relevant quantity.

**FIG. 3.**
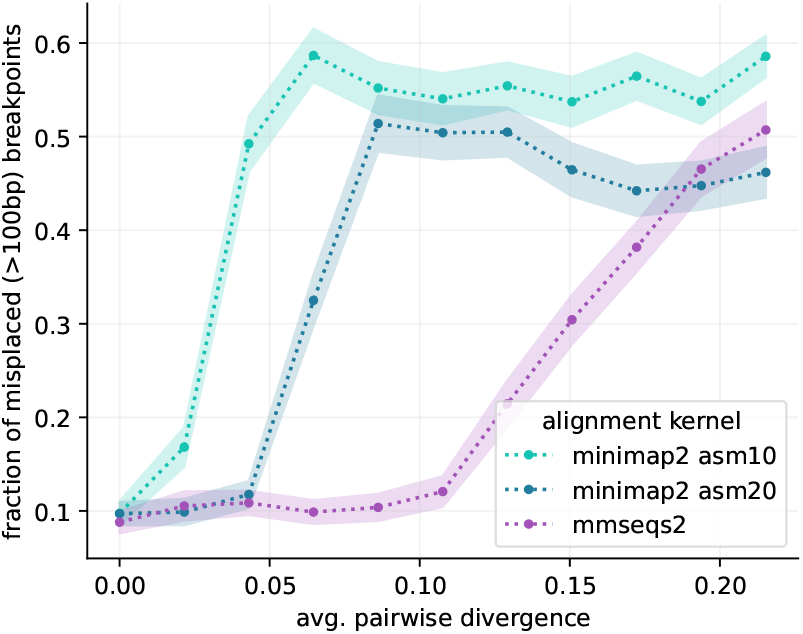
Accuracy against synthetic data. We generate artificial data with varying degree of sequence divergence, and compare the real underlying pangenome graph with the one reconstructed by *PanGraph*, for three different alignment kernels *minimap2* with *asm10* or *asm20* option, and *mmseqs2*. In each comparison we evaluate the misplacement of breakpoints that we can pair on the two graphs within 1kbp. The plot displays the fraction of breakpoints that have misplacement greater than the standard *PanGraph* precision threshold of 100bp, as a function of average pairwise sequence divergence. Line and shaded area represents mean and standard deviation over 25 repetitions. *mmseqs2* maintains accuracy at higher divergence, at the cost of higher computational time.

### B. Validation on real data

We additionally validated PanGraph on genomes from natural populations sampled from RefSeq (O’Leary *et al*., 2016), focusing on the properties of the resulting pangenome graphs as a function of dataset size and diversity.

#### Dataset characterization

We downloaded from RefSeq (O’Leary *et al*., 2016) completely assembled chromosomes from 5 different bacterial species: *Escherichia coli* (EC), *Klebsiella pneumoniae* (KP), *Helicobacter pylori* (HP), *Mycobacterium tuberculosis* (MT) and *Prochlorococcus marinus* (PM). These data had been previously analyzed using PanX (Ding *et al*., 2018), which allowed us to estimate the size of the pangenome and core genome, and the average pairwise divergence on core genes (cf. Table I and SI Section IIIA).

**TABLE I.** Dataset properties. Columns represent: number of isolates (N.), average chromosome length (⟨*L*_gen_⟩), total pangenome length (*L*_pang_), total core genome length (*L*_core_), average pairwise divergence of core genes (⟨*d*_core_⟩).

| Species | N. | $\langle L_{\text{gen}} \rangle$<br>[Mbp] | $L_{\text{pang}}$<br>[Mbp] | $L_{\text{core}}$<br>[Mbp] | $\langle d_{\text{core}} \rangle$ |
| --- | --- | --- | --- | --- | --- |
| *E. coli* | 307 | 5.0 | 17.7 | 0.7 | 1.6% |
| *K. pneumoniae* | 109 | 5.3 | 13.0 | 2.3 | 0.8% |
| *H. pylori* | 85 | 1.6 | 1.9 | 0.6 | 4.2% |
| *M. tuberculosis* | 51 | 4.4 | 4.1 | 2.4 | 0.03% |
| *P. marinus* | 10 | 1.8 | 3.3 | 0.7 | 26.9% |

Among these 5 data sets, the *E. coli, K. pneumoniae, M. tuberculosis* genomes have core genome diversity below the thresholds of the minimap2 aligner, while *H. pylori* is diverse enough that we don’t expect minimap2 to find all relevant matches, while mmseq2 should process this set without problems. *P. marinus*, on the other hand, is very diverse with an average core genome diversity of 26% and is beyond what we expect PanGraph to handle.

#### Benchmark on real data

Using *PanGraph*, we built multiple pangenome graphs for each species in the dataset (cf. SI Section IIIB and IIIC). These differ by the alignment kernel used (*minimap2* with *asm10* or *asm20* option, and *mmseqs2*) or by the value of the pseudo-energy parameters *α, β* from eq. (1). Different alignment kernels are expected to reach different accuracy on datasets with different diversities (cf. Fig. 3). Moreover the use of null energy parameters (*α* = *β* = 0) is expected to remove the threshold divergence of 10% associated with the standard values of parameters (*α* = 100, *β* = 10) at the cost of more fragmented pangenome graphs.

*PanGraph* can build pangenome graphs comprising hundreds of isolates in a few hours (5h for 307 E. Coli isolates with *minimap2* kernel, cf. Fig. 4A). Use of *mmseqs2* consistently requires longer computational time, but provides higher sensitivity when merging highly diverged sequences.

**FIG. 4.**
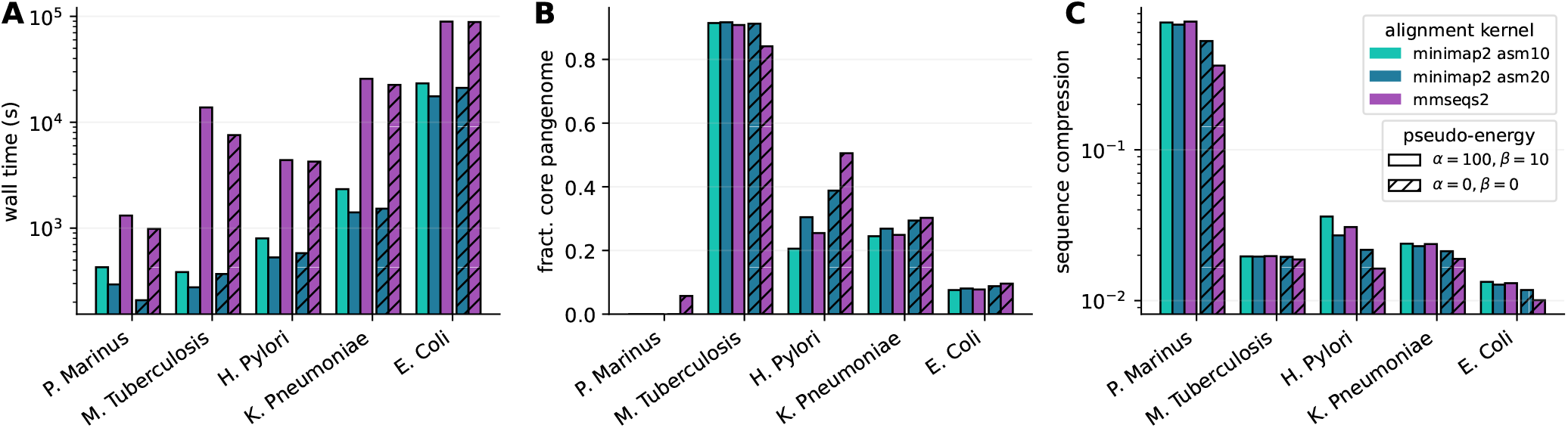
Benchmark on real data. We build pangenome graphs from fully-assembled chromosomes from 5 different bacterial species. For each species we build graphs with three different alignment kernel options (*minimap2* with *asm10* or *asm20* option and *mmseqs2*) and two different settings for the pseudo-energy parameters *α, β* (standard or null values). **A**: *PanGraph* wall-time when run in parallel on 8 cores. **B**: fraction of core pangenome in the pangenome graph. **C**: sequence compression, defined as the ratio between the pangenome graph size and the cumulative size of all the sequences contained in the graph.

Fig. 4B-C and SI Fig. 4 quantify the size of the core genome, i.e. the sum of all pancontigs present once in every genome, identified by PanGraph and the compression of the dataset, i.e. the ratio for the sum of the lengths of all pancontigs and the sum of the lengths of all input genomes, for different dataset and the different parameters. As expected, the very homogenous *M. tuberculosis* dataset has a large core genome with and a compression ratio that is almost equal to the inverse number of input genomes, independent of the aligner or parameters used. *E. coli* and *K. pneumoniae* compress similarly well, but their core genome is a much smaller fraction of the pangenome. Using the more sensitive aligner mm-seqs2 and relaxing the merger parameters leads to some additional compression, presumably due to merging of diverged paralogs.

The *H. pylori* dataset sits at the limit of what can be accurately merged with the *minimap2* kernel. In this case the use of *mmseqs2* and null energy parameters increases core pangenome fraction, decreases the total pangenome size and and does not compromise the fragmentation of the graph (cf. Fig. 4B-C and SI Fig. 4). The divergence of the *P. marinus* dataset sits instead beyond the capabilities of *Pangraph*. In this case no alignment kernel can reach satisfactory sequence compression, and only *mmseqs2* combined with null energy parameters is able to retrieve a few core pancontigs.

For species other than *P. marinus*, the pangraphs contain thousands to tens of thousands of pancontigs, and 50% of the pangenome sequence is contained in blocks spanning several kbps (see SI Fig. 4). For these species the use of null energy parameters slightly increases merging and sequence compression, at the cost of slightly more fragmented graphs (cf. SI Section IIIC and SI Fig. 4). Overall this benchmark showcases the capabilities and limits of *PanGraph*, and demonstrates the value of adapting the pseudo-energy parameter and choice of alignment kernel to the expected diversity of the dataset considered.

#### Scaling with dataset size

We then explored how the properties of the pangraph scale with the dataset size. To do so, we build pangenome graphs from randomized subsets of isolates of increasing size from the *E. coli* dataset. Results are reported in Fig. 5.

**FIG. 5.**
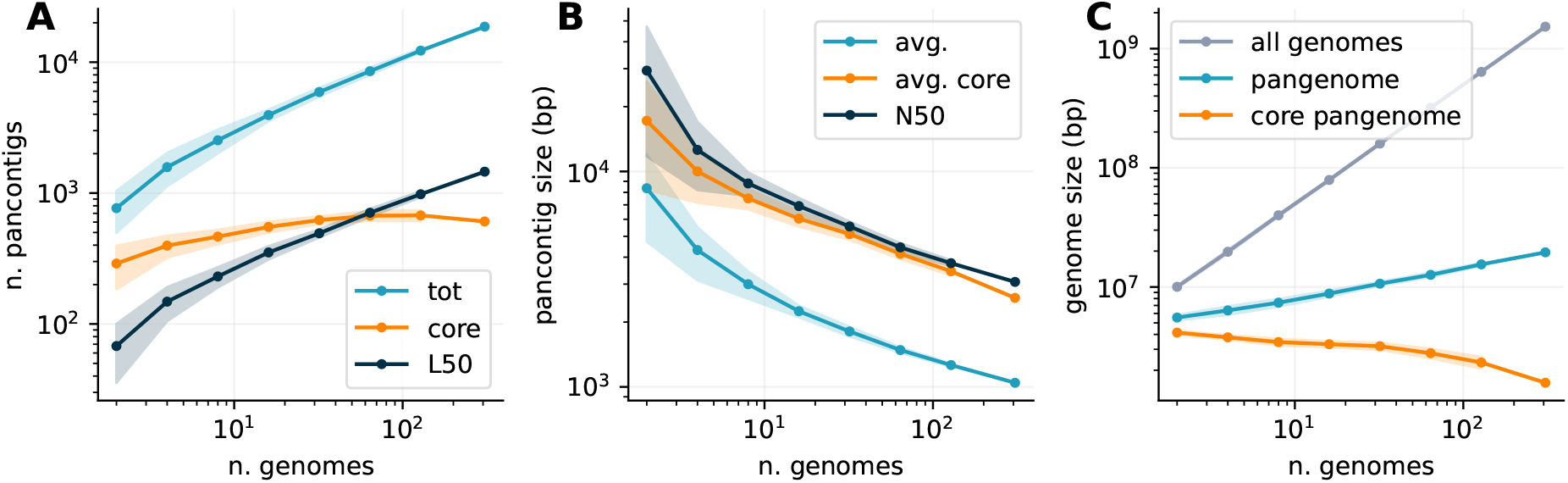
Pangenome graph properties vs. dataset size. We build pangenome graphs with increasing number of isolates from the E. Coli dataset and measure the scaling of different properties of the graphs. Graphs were built using *minimap2* alignment kernel with *asm20* option. Lines and shaded areas represent mean and standard deviation over 10 different repetitions on random subsets of isolates, except for the final point indicating the full graph (307 isolates). **A**: n. of pancontigs in the graph. We count the total number of pacontigs (blue), the number of core pancontigs (orange) and the minimum number of pancontigs that contain more than 50% of the pangenome (L50, black). **B**: average size of pancontigs (blue), of only core pancontings (orange), and size of the smallest panconting in the minimal set that spans 50% of the pangenome (N50, black). **C**: cumulative size of all genomes in the pangenome graph (gray), total pangenome size (blue) and size of core pangenome (orange).

The number and size of pancontigs scale sub-linearly with the number of isolates, with more than 50% of the pangenome being included in the 10% longest pancontigs, and core pancontigs having higher-than-average size. Adding more strains does not generate excessive fragmentation of these pancontigs, and their size remains of several kbps even as hundreds of strains are added. The size of the core genome does not decrease significantly with the addition of new strains, while the total pangenome size remains orders of magnitude smaller than the total size of all genomes included in the graph.

## III. GRAPH MARGINALIZATION

The interpretation of a pangenome graph containing hundreds of strains can be challenging. To this end, it is often informative to inspect simpler sub-graphs comprising only a subset of isolates. However building many such sub-graphs is computationally intensive. To facilitate this task *PanGraph* provides the `marginalize` command, that can be used to project a large pangenome graph on a small subset of strains, removing all the other paths and merging transitive pancontigs. This operation is computationally much cheaper than building a new graph for the subset of strains considered.

In addition to being a useful operation, marginalization allows us to quantify the consistency and robustness of pangraph construction on real data, where the ground truth is unknown. To verify that marginalized graphs are compatible with newly-built graphs, we built a pangenome graph for 50 randomly selected strains from the *K. pneumoniae* dataset. We then picked 50 random pairs of strains from the same set and built two graphs for each pair: the graph obtained by marginalizing the complete graph on the pair (i.e. marginalized graph in Fig. 6A top), and the graph built directly from the pair (i.e. pairwise graph in Fig. 6A top). In order to compare both graphs, we compute the partition they generated on the genome of the isolates they include. Each genome is partitioned in pancontigs that are either shared on the pair, or private to one isolate (cf. Fig. 6A bottom). We can classify the segments in the intersection of the two partitions in different categories, depending on whether the two partitions agree on whether the segment is shared or private. Moreover segments on which the two partitions agree are further sub-divided depending on whether they are private or shared on both partitions.

**FIG. 6.**
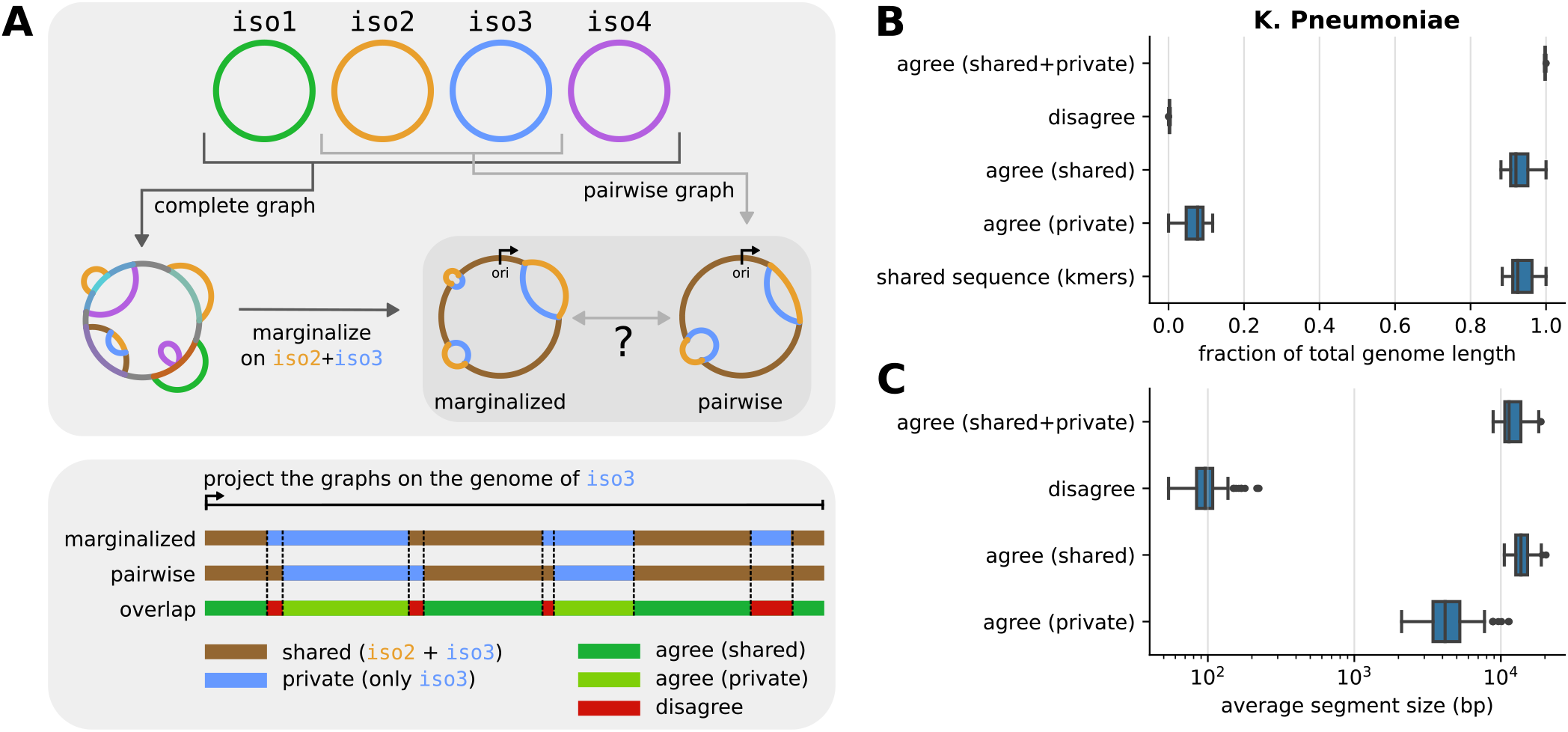
Test of graph marginalization. **A**: we built a pangenome graph from 50 randomly chosen strains from the *K. pneumoniae* dataset. We then randomly picked 50 pairs of strains. For each pair we compared the pangenome graph obtained by marginalizing the complete graph on the pair of strain, and the one obtained by building a new graph for the pair (top). The comparison is done by considering that each graph partitions a genome in shared and private segments. By combining the partitions generated by the marginalized and pairwise graphs we categorize segments in three categories, depending on whether the two partitions agree or not, and if they agree depending on whether segments are shared or private. All graphs were built using *minimap2* alignment kernel with *asm20* option and default value for the energy parameters. **B**: distribution of the average fraction of the genome covered by segments of each category, over the 50 pairs considered (2 entries per pair). The last line represents the distribution of shared sequence, approximated using the fraction of shared k-mers corrected using sequence divergence as described in the main text. **C**: distribution of average segment lengths for each category over the 50 pairs considered (2 entries per pair).

The compatibility between the two graphs requires segments on which the two partitions disagree to be few and short. Indeed, we verified that over the pairs we picked, these segments cover a very small fraction of the genome (*<* 1%) and have average size compatible with the default 100 bp precision threshold of *PanGraph* (cf. Fig. 6B-C). Conversely, segments on which the partitions agree have average size of several kbps. The fact that the two partitions almost completely agree cannot be explained simply by the fact that most of the genome is shared, since segments that are shared on both graphs cover on average only 85% of the sequence. We confirmed these results using the fraction of shared k-mers, using an approach inspired by PopPUNK (Lees *et al*., 2019). Namely, we approximate the fraction of homologous sequence between any two pairs using the fraction of shared k-mers (*k* = 21) divided by (1 *− d*)^*k*^, where *d* is the average pairwise divergence on core genes for the pair considered. This divisor corrects for k-mers on homologous sequence that are not shared due to mutations. The resulting distribution of shared sequence is in very good agreement with the fraction of segments that are shared on both graphs (cf. Fig. 6B), suggesting that homologous sequences are correctly merged on the complete, marginalized and pairwise graphs.

We performed the same test on species from the other datasets, obtaining similar results on all species whose divergence is compatible with *PanGraph* capabilities (cf. SI Section IV and SI Fig. 5).

## IV. DISCUSSION

While single nucleotide differences in the core genome are straightforward to analyze with existing tools, these analyses miss the great majority of genetic diversity. The ability to rapidly align large sets of complete genomes of a bacterial species is crucial for the investigation of the processes governing microbial evolution. We developed *PanGraph* as a tool that can address this need, by being able to capture the structural and nucleotide diversity in both the core and accessory genome in a scalable way.

In our analysis we demonstrated the capabilities and limits of this tool, both on synthetic and real sequences. The efficient implementation of *PanGraph* allows it to create pangenome graphs containing hundreds of isolates in a few hours on a 8-core machine. The size of pancontigs and the fraction of core sequence have good scaling properties with the number of isolates in the graph, indicating that *PanGraph* is able to successfully capture pangenome properties. By construction *PanGraph* operates in a gene-agnostic way, being only based on sequence homology, and is thus robust to gene annotation errors. One of the main limitations is the diversity of the input sequences. With the default *minimap2* alignment kernel, *PanGraph* is able to correctly merge genomes with up to 5% average divergence. Sequences with higher divergence (up to 10-15%) can be merged using the *mmseqs2* alignment kernel, and tuning the *α, β* energy parameters (cf. eq. (1)), trading off speed for sensitivity. For more diverged data sets, homology detection tools that use protein sequences would likely be necessary.

We also provide the ability to quickly marginalize big graphs on a subset of strains, obtaining simpler graphs that can more easily be explored. Graphs can also be exported in different formats for further analysis and visualization.

The applications of *PanGraph* are not limited to whole bacterial chromosomes. Plasmids and other Mobile Genetic Elements (MGEs) play a fundamental role in microbial evolution (Frost *et al*., 2005; Haudiquet *et al*., 2022), including in the spread of antibiotic resistance (Van der Zee *et al*., 2018). MGEs often display a wide variety of sequence and structural diversity. Being able to capture and represent this diversity is key to understanding their evolution, particularly on epidemiologically relevant timescales where very few SNPs accumulate but large-scale structural changes are frequent (Noll *et al*., 2018; Sheppard *et al*., 2016). In this sense, the pangenome graph is a naturally powerful representation, and *PanGraph* can be used to explore this diversity (Shaw and Neher, 2022). Some example applications on plasmids are available in *PanGraph*’s documentation^2^.

We hope that *PanGraph* will prove a valuable tool, with the potential to spur new insights into microbial diversity and the processes by which bacteria adapt and change.

## ACKNOWLEDGEMENT

We are grateful to Boris Shraiman for stimulating discussions. This study was funded by the University of Basel and the NSF. L.P.S. is a Sir Henry Wellcome Postdoctoral Fellow funded by Wellcome (Grant 220422/Z/20/Z).

## Footnotes

1 Instructions on how to run the analysis scripts are available at https://github.com/neherlab/pangraph/blob/master/script/README.md

2 See the tutorial section in https://neherlab.github.io/pangraph

