## Supplementary information for "PanGraph: scalable bacterial pan-genome graph construction"

### PanGraph: scalable bacterial pan-genome graph construction
### Supplementary Information

Nicholas Noll,[1,*] Marco Molari,[2,3,*] Liam P. Shaw,[4] and Richard A. Neher[2,3]
[1]*Kavli Institute for Theoretical Physics, University of California, Santa Barbara*
[2]*Swiss Institute of Bioinformatics, Basel, Switzerland*
[3]*Biozentrum, University of Basel, Basel, Switzerland*
[4]*Department of Biology, University of Oxford, Oxford, UK*

#### I. GENERATION OF SYNTHETIC DATA

As a first test we benchmark the performances and accuracy of *PanGraph* on synthetic data, obtained by simulating the evolution of a population in which individuals accumulate mutations, can loose part of the sequence from deletions, or gain new genetic material via Horizontal Gene Transfer (HGT) from other members of the population.
In the simulation a population of size $N$ is evolved for $T$ generations using a Wright-Fisher model (Hudson, 2002). Each individual in the population has a circular genome of initial size $L$. At each generation single-nucleotide mutations occur at a rate $\mu$ per position. Deletions can occur at a rate $d$ per genome per generation. When a deletion is suggested on isolate $n$, having a genome length $L_n$, a new desired length $L'_n$ is extracted from a Gaussian distribution with mean $L$ and variance $\sigma^2$. If $\Delta = L'_n - L_n < 0$ then a random chunk of length $|\Delta|$ is removed from the genome, reducing it to size $L'_n$. If $\Delta > 0$ no deletion is performed. This ensures that the length distribution of genomes in the population remains close to the desired length $L$, with $\sigma^2$ being a proxy for the variance. HGT occurs at a rate $h$ per genome per generation. Similarly to deletions, when a HGT event is suggested a new desired length $L'_n$ is extracted from the same Gaussian distribution, and this time the event is performed only if $\Delta = L'_n - L_n > 0$. In this case a random chunk of sequence of length $\Delta$ is extracted from a random individual in the parent population (possibly the parent of isolate $n$ itself) and inserted in a random position of the genome of isolate $n$. Finally, inversions occur at a rate $i$ per genome per generation. When an inversion is proposed a random length $\Delta$ is chosen in the same way, and a random chunk of length $|\Delta|$ is inverted. Standard values of the simulation parameters are reported in Table I.

| Parameter | Description | Standard Value |
|---|---|---|
| $N$ | population size | 100 |
| $T$ | n. of simulated generations | 50 |
| $L$ | average genome size | 50 000 [bp] |
| $\sigma$ | s.t.d. of genome size | $L/10$ [bp] |
| $\mu$ | mutation rate | 0.005 [1/bp · generation] |
| $d$ | deletion rate | 0.05 [1/generation] |
| $i$ | inversion rate | 0.01 [1/generation] |
| $h$ | HGT rate | 0.1 [1/generation] |

TABLE I **Simulation parameters**. Description of all simulation parameters used for the generation of synthetic data. Unless otherwise specified, the standard value is used.

At the end of the simulation the resulting mosaic genomes are collected, along with the real underlying pangenome graph that can be used as the ground truth against which to check the performances of *PanGraph*. The code to perform these simulations is shipped with *PanGraph* in the `generate` command.

#### II. BENCHMARK OF PANGRAPH ON SYNTHETIC DATA

We test the performance of *PanGraph* on synthetic data, focusing on two aspects: the computational performances of the algorithm (time and memory requirements) as a function of the dataset size, and the accuracy of the algorithm

* These two authors contributed equally

in reconstructing the real pangenome graph of the population as a function of sequence divergence.

##### A. Computational performance

To test the computational performances of *PanGraph* we generated data using the model described in the previous section, with standard value of the parameters but varying the average genome length $L = [1, 5, 10, 50, 100, 500]$ kbp and the population size $N = [10, 20, 50, 100, 200, 500, 1000]$. For each pair of $N, L$ values we generated 50 different datasets. On each dataset we ran the *PanGraph* `build` command, using *minimap2* as alignment kernel with *asm20* option. We used standard value for the energy parameters $\alpha = 100$, $\beta = 10$. Runs were performed using 8 cores. For each run we measured the wall-time of the command, the maximum memory requirements (maximum resident size) and the average cpu percent. Results are displayed in Fig. 1. Thanks to the guide tree architecture of *PanGraph* the run time scales almost linearly with the number of isolates.

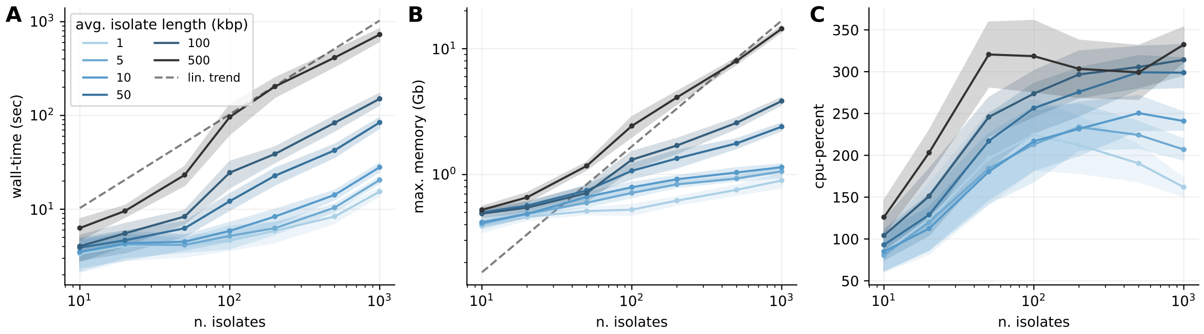


FIG. 1 **Computational performance of PanGraph algorithm on synthetic data, as a function of dataset size**. We run PanGraph on artificially generated datasets consisting of a variable number of isolates with genomes of variable length (colorscale). For each condition we display the average and standard deviation over 50 different runs of: (A) algorithm wall-time, (B) maximum memory requirement and (C) average percentage of cpu used on a total of 8 cores.

##### B. Accuracy in reconstructing the pangenome graph

To test the accuracy of *PanGraph* in reconstructing the pangenome graph of a set of isolates we generated artificial data using the procedure described in Section I. For each simulation we obtain both a set of genome sequences and the underlying real pangenome graph. This was done using standard value of the parameters, but varying the rate of HGT in the interval $h = [0.01, 1]$ and the mutation rate $\mu = [0, 0.01]$. For each $(h, \mu)$ pair 25 different sets of data were generated.
On each set of data we executed *PanGraph* with values for the energy parameters $\alpha = 0$ and $\beta = 0$, and with three different alignment kernels:

- *minimap2* with option *asm10*,
- *minimap2* with option *asm20*,
- *mmseqs2*,

thus obtaining three different pangenome graphs per dataset.
To link the mutation rate parameter $\mu$ with the average pairwise SNPs distance $\langle d \rangle$ between homologous segments in the population we evaluated the average pairwise distance for every pancontig in the pangenome graph with depth greater than one. For each graph we then perform the weighted average of these divergences, using as weight the length of the pancontig. Finally, for every pair of parameters $(\mu, h)$ we average these numbers over the 25 different simulations. Results are displayed in Fig. 2.
For high values of $\mu$, the average pairwise divergence of pancontigs is not influenced by the rate of HGT $h$, but depends strongly on the choice of alignment kernel. With the *asm20* option, *minimap2* is able to correctly merge genomes with average pairwise divergence $\langle d \rangle \sim 7\%$ in our simulations, while *mmseqs2* reaches $\langle d \rangle \sim 10\%$ at the expense of higher processing time. Notice that these values are lower than the threshold sensitivity of these aligners,

which are expected to find matches up to respectively around 10%, 20% and 30% sequence divergence. This is due to the fact that homologous sequences in our simulations are generated by an evolutionary process, and they can have a wide distribution of diversities and hierarchical population structure. Inability to merge sequences above a threshold divergence in a single panconting is expected to result in multiple pancontigs containing sub-clades with higher similarity, and average diversity that is appreciably lower than the aligner threshold.
By performing a linear fit on the datapoints with $\mu \leq 0.002$ we are able to recover the conversion factor between the mutation rate and average pairwise divergence of pancontigs in our simulations: $\langle d \rangle \sim 20.8\,\mu$.

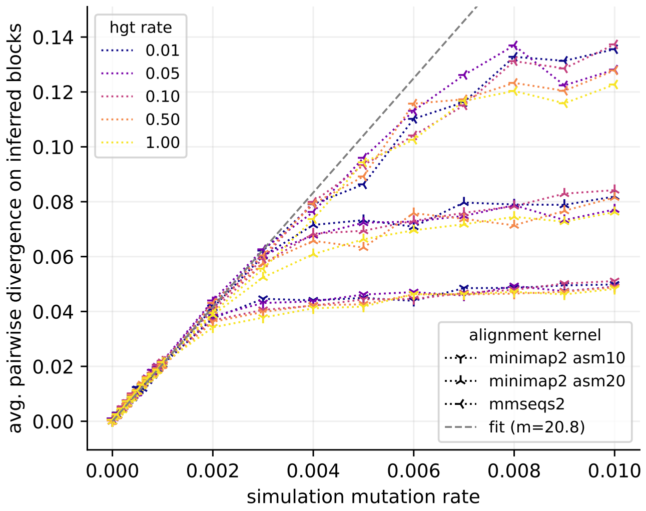


FIG. 2 **mutation rate vs divergence on synthetic data**. We compare the mutation rate $\mu$ of our simulations with the average pairwise divergence $\langle d \rangle$ of sequences in pancontigs of the resulting pangenome graphs. This is done for different values of the rate of HGT $h$ and for three different alignment kernels. As expected, the divergence is only marginally influenced by the HGT rate, but the saturating value of the divergence depends strongly on the choice of alignment kernel, with *mmseqs2* being able to merge sequences with average divergence higher than 10%. To find the relationship between the mutation rate $\mu$ and the average pairwise divergence of the sequences we peform a linear fit on data points for $\mu \leq 0.002$. This provides the conversion factor $\langle d \rangle \sim 20.8\,\mu$.

Once this link has been established we group simulations by the value of $h$ and measure accuracy by comparing the reconstructed pangenome graphs with the ground truth provided by our simulations. In particular, for each isolate we consider the breakpoints between different pancontigs that tile the genome, and measure the displacement between the real position of these breakpoints and the position reconstructed by pangraph.
To evaluate this displacement it is first necessary to link breakpoints on the real and reconstructed pangenome graph. We formulate this problem as an assignment problem, and solve it numerically using the Hungarian method (Kuhn, 1955). For each isolate we call $b_i$ for $i = 1, \ldots, I$ the pancontig breakpoints on the real pangenome graph, and $b_j$ for $j = 1, \ldots, J$ the ones on the reconstructed pangenome graph.[1] We define a cost matrix $D_{ij} = d(b_i, b_j)$ whose elements are distances on the genome of pairs of breakpoints from the two graphs. We then numerically solve the assignment problem on this matrix, and obtain a set of $K = \min\{I, J\}$ pairings between breakpoints such that the distance between the pairs is minimal and all the breakpoints of the graph with minimum number of breakpoints have been paired. The average distance between breakpoints in these pairs is the average breakpoint distance of the graph.
For each simulation we obtain a number of average breakpoint distances equal to the number of isolates in the simulation. In Fig. 3 we plot the cumulative distribution of these distances, with a cutoff at 1kbp. We stratify simulations according to the average pairwise divergence of pancontigs inferred from the value of $\mu$. Each panel corresponds to a different alignment kernel. As the sequence diversity increases we observe a clear transition. For low diversity most of the breakpoints are inferred to be only a few bp away from their real position, while for highly diverged sequences the position of breakpoints is not precisely inferred and can be hundreds of bp away from their

[1] To avoid artifacts generated by the default threshold distance $d = 100$ bp of *PanGraph*, if any pair of breakpoints on the same graph sits at a distance smaller than this threshold, we remove one of them.

real position. For different alignment kernels this transition occurs at different values of the divergence, with *mmseqs2* being the most accurate, as can be visualize from Fig. 3 in the main text.

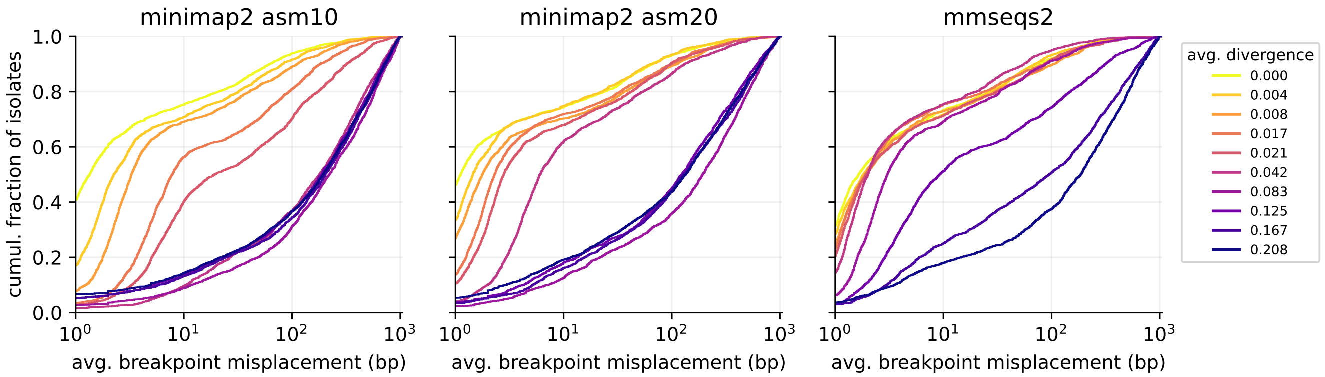


FIG. 3 **Accuracy of PanGraph algorithm on synthetic data**. For each of our generated datasets we evaluate the average displacement of inferred pancontig breakpoints. The figure represents the cumulative distribution of average misplacements, stratified by the average pairwise divergence of sequences in pancontigs (see colorbar on the right). Each panel corresponds to a different alignment kernel used to infer the pangenome graph (see panel title). As divergence increases we observe a transition in the precision of breakpoint position estimation. At low divergence the position of most breakpoints is correctly estimated withing a few bps, while at high divergence most breakpoints can be misplaced by multiple hundreds of bps. The transition value for the divergence depends on the alignment kernel used, with *mmseqs2* having the highest accuracy at the cost of higher computational time.

#### III. BENCHMARK OF PANGRAPH ON REAL DATA

We benchmark *PanGraph* on real data, focusing on the computational performances and the properties of the resulting pangenome graphs.

##### A. Dataset selection

We downloaded sets of annotated and complete chromosomes from RefSeq (O'Leary *et al.*, 2016) for 5 different bacterial species: *Klebsiella pneumoniae* (KP), *Helicobacter pylori* (HP), *Prochlorococcus marinus* (PM), *Mycobacterium tuberculosis* (MT) and *Escherichia coli* (EC). In Table I in the main text we report some summary statistics for these datasets. The number of isolates per species ranges from several hundreds for EC to 10 for PM. These data were previously analyzed using PanX (Ding *et al.*, 2018), a pipeline for the analysis of bacterial pangenome. The pipeline takes as input a set of annotated genomes, and it clusters and aligns homologous genes. This allowed us to identify core genes, and evaluate their average pairwise sequence divergence $\langle d_{\mathrm{core}} \rangle$. We also evaluated the average genome length $\langle L_{\mathrm{gen}} \rangle$, the total pangenome length $L_{\mathrm{pang}}$ and core genome length $L_{\mathrm{core}}$ (see Table I in the main text). EC is the dataset with the highest number of isolates (307) and is also the one with the biggest pangenome. MT is the species with the smallest sequence divergence ($\langle d_{\mathrm{core}} \rangle \sim 0.03\%$) and the biggest core genome fraction. The two datasets with highest sequence divergence are HP ($\langle d_{\mathrm{core}} \rangle \sim 4\%$) and PM ($\langle d_{\mathrm{core}} \rangle \sim 24\%$). For HP the average sequence divergence sits on the limit of what *PanGraph* is able to accurately merge when using the *minimap2* kernel (cf. Fig. 3), but is merged correctly by the *mmseqs2* kernel. The sequence divergence of PM sits instead beyond the capabilities of all alignment kernels.

##### B. Pangenome graph construction

We built pangenome graphs using five different options for the alignment kernel:

- kernel *minimap2* with option *asm10* and default values for the energy parameters $\alpha = 100$, $\beta = 10$.
- kernel *minimap2* with option *asm20* and default values for the energy parameters $\alpha = 100$, $\beta = 10$.

- kernel *mmseqs2* and default values for the energy parameters $\alpha = 100$, $\beta = 10$.
- kernel *minimap2* with option *asm10* and null energy parameters $\alpha = 0$, $\beta = 0$.
- kernel *mmseqs2* and null energy parameters $\alpha = 0$, $\beta = 0$.

From the results described in the previous section (see Fig. 3) we expect *mmseqs2* to be able to merge sequences with higher divergence than *minimap2*. Moreover from the definition of the pseudo-energy eq. (1) in the main text, the value of the energy parameters $\beta$ defines an upper threshold for sequence divergence $d \sim 1/\beta$. Homologous sequences with divergence higher than this value will result in positive pseudo-energy and will not be merged in a single pancontig. For $\beta = 10$ the critical divergence threshold is $d \sim 10\%$. Executing pangraph with the option $\alpha = 0$, $\beta = 0$ will result in the merging of more diverged sequences, at the cost of a more fragmented pangenome graph containing a higher number of shorter pancontigs.

##### C. Benchmark results

The results of the benchmark are displayed in Fig. 4. In each panel data are divided by species, with colors and texture identifying the five possible choices for the alignment kernel.
Execution time (cf. panel B) depends on the dataset size and ranges from few minutes for PM to several hours for EC. Aligning sequences with *mmseqs2* consistently requires more time than *minimap2*. The time required to build a pangenome graph for the 307 chromosomes of EC is only 5h for *minimap2*, and around 24h for *mmseqs2*. The latter also has a higher minimum memory requirement (panel C, around 8 Gb when running on 8 cores) but this is comparable to *minimap2* when aligning several tens of isolates.
We quantify the properties of pancontigs by looking at three main statistics: their total number (panel D) the size of the minimal set of pancontigs that includes at least 50% of all the graph sequence (L50 statistics, panel E) and the length of the shortest pancontig in this minimal set (N50 statistics, panel F). As expected, the use of null energy parameters results on average in a more fragmented graph, having more and shorter pancontigs. In general the pangenome graphs of our datasets have around $10^3$ / $10^4$ pancontigs, with 50% of the sequence being contained in around 10% of them, having N50 length of around 1-10 kbp.
The total pangenome length (panel G) depends both on the number of isolates and the ability of the alignment kernel to merge homologous sequences. When setting the energy parameters to zero more diverged sequences are merged into the same pancontigs and the total pangenome graph size is consistently smaller. The total graph size is in general much smaller than the cumulative size of the genomes it contains (panel I), with compression up to 1-2% of the total size for graphs with a high number of isolates (EC) or with small accessory genome (MT). The fraction of core sequence (panel H) is consistent with the species expectation, with MT having the highest core fraction.

Two species in our datasets hit the limits of the capabilities of *PanGraph*: HP and PM.
HP contains some homologous sequences with divergence higher than the default threshold $d = 1/\beta \sim 10\%$. Removing this threshold by using null energy parameters results in a graph with fewer pancontigs, smaller L50, greater fraction of core pangenome and higher level of sequence compression. These are all indications of more complete merging of homologous sequences. This effect is more marked for *mmseqs2* than for *minimap2*, in agreement with their different divergence limit (cf. Fig. 3). The HP datasets represents an example in which relaxing the default divergence threshold of pangraph and using a more accurate alignment kernel results in a more consistent pangenome graph.
Conversely, the PM dataset contains highly diverged isolates, whose divergence go beyond the capabilities of both alignment kernels ($\langle d_{\text{core}} \rangle \sim 27\%$, cf. Table I in the main text). Only *mmseqs2* manages to partially merge some of the least diverged core sequences. This results in a small core genome fraction and poor sequence compression. In its current state *PanGraph* cannot meaningfully merge genomes with such high divergence into a pangenome graph.

#### IV. GRAPH MARGINALIZATION

Given a pangenome graph containing many different genomes, *PanGraph* is capable of marginalizing it on a subset of strains with the `marginalize` command. Marginalization removes undesired paths and merges any remaining transitive pancontigs, i.e. pancontigs that are co-linear in all of the remaining paths. The resulting graph should be equivalent to the one obtained by directly building a graph on the selected subset of strains, with the added advantage that marginalization is a fast operation. This makes it more convenient to build a pangenome graph for the full set of

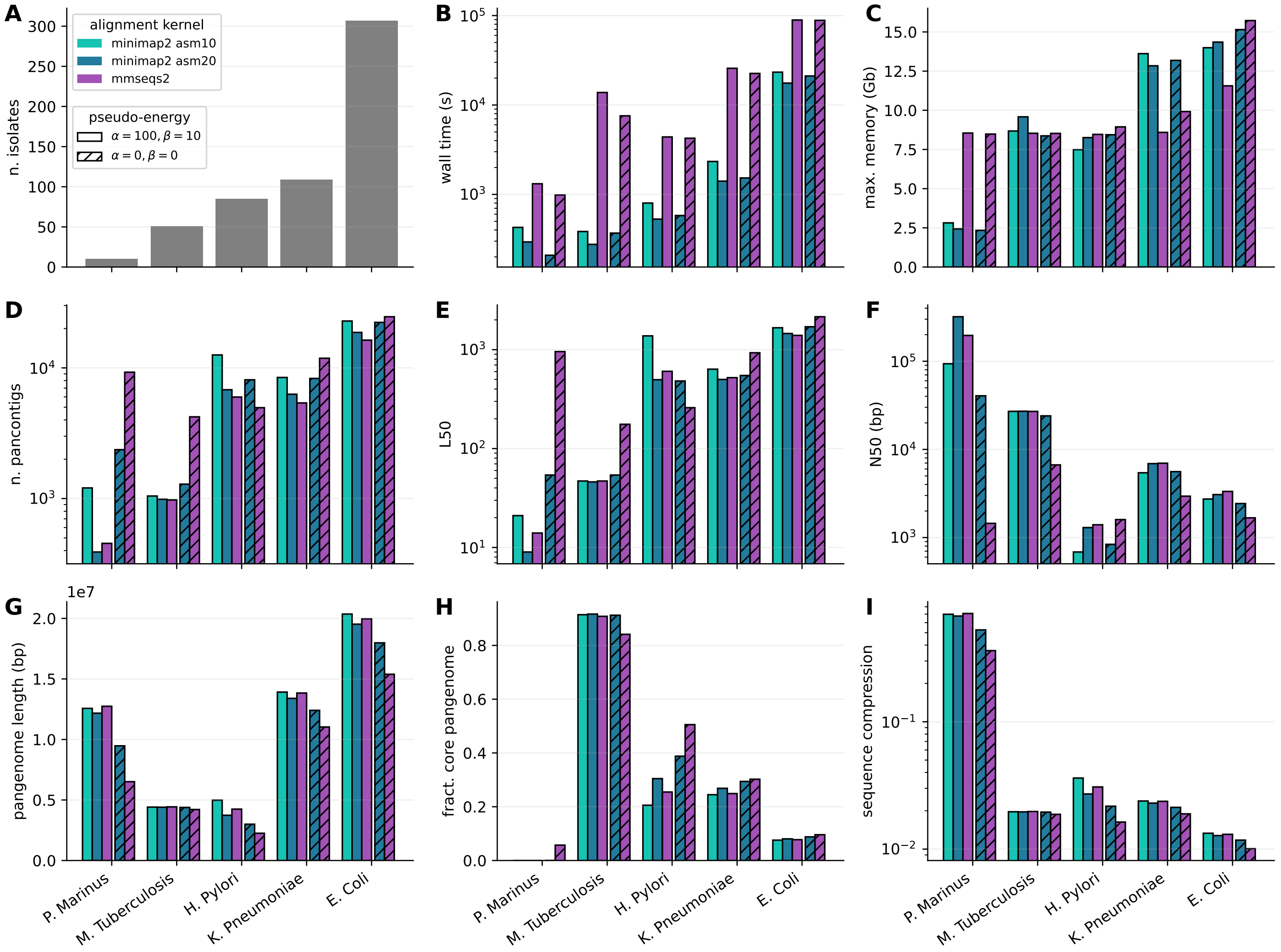


FIG. 4 **Benchmark on real data**. We run *PanGraph* on datasets consisting of 5 different bacterial species, and using 5 different options for the alignment kernel as described in the text. Runs were performed using 8 cores. **A**: number of isolates for each species. **B**: algorithm wall-time. **C**: maximum memory usage. **D**: total number of pancontigs in the pangenome graph. **E**: minimum number of pancontigs that can add up to 50% of the total pangenome graph length. **F**: threshold length such that the cumulative size of all pancontigs longer than this threshold is more than 50% of the full pangenome graph. **G**: total length of the pangenome graph (sum of all pancontig lengths). **H**: fraction of the pangenome graph that is composed of core pancontigs, i.e. pancontigs that occur exactly once per isolate. **I**: ratio between the length of the pangenome graph and the total length of all genomes included in the graph.

isolates considered, and then marginalizing it multiple times on interesting subsets, rather than building a new graph for each subset.

We used genomes from the dataset described in the previous section to verify that marginalized graphs are compatible with graphs built directly from subset of strains. For each species we selected 50 different isolates (10 for PM) and built pangenome graphs using standard parameter values and the *minimap2* kernel with *asm20* option. For each species we then randomly picked 50 different pairs of isolates (all 45 for PM).[2] For each pair we then built pairwise graphs using the same parameters, and compared them to the corresponding marginalized graph, obtained by projecting the full pangenome graph for the 50 isolates on the selected pair (cf. Fig. 6A, main text).
The comparison was performed by considering that each graph partitions a genome into pancontigs. Each pancontig can either be shared with the other member of the pair, or can be private to the isolate (cf. Fig. 6A, main text). For each isolate we consider the two partitions defined by the marginalized and pairwise graphs, and generate their

[2] The list of selected accession numbers and selected pairs for each species is available on the repository: `https://github.com/neherlab/pangraph/blob/master/script/config/projection_strains.json`

intersection. Each block in this intersection can belong to two different categories, depending on whether the two partitions agree or disagree on whether the block is shared. Moreover, we can separate blocks on which we find agreement in two further categories: blocks that are shared and blocks that are private on both graphs.

For each pair of genomes and each type of block we evaluate the fraction of the genome that they cover and their average size. In Fig. 5 we display the distribution of these quantities across all pairs for the different species considered and the block types described above.

In all species except HP and PM we find very good agreement between pairwise and marginalized graphs: segments on which the graphs *disagree* represent only a very minor fraction of the genome (much smaller than the fraction of shared or private blocks), and have size compatible with the default block length threshold of *PanGraph* (100 bp). As described in the previous section, the divergence of the HP dataset sits on the limit of what the *minimap2* alignment kernel can successfully handle. In this case the results of merging different graphs together can depend on the order of merging, and pairwise and marginalized graphs can potentially show some disagreement. In this case the disagreement remains minor and only extends to few percents of the genome size, with disagreement blocks having size only slightly bigger than the 100 bp threshold of *PanGraph*. PM represents instead an example of a dataset with divergence much beyond the limit of the alignment kernel ($\langle d_{\mathrm{core}} \rangle \sim 27\%$). As a consequence, highly diverged homologous sequences are not merged, and the great majority of segments are private and have large size. Results also show a strong order-dependence, with some *disagreement* blocks having size of 10s of kbp.

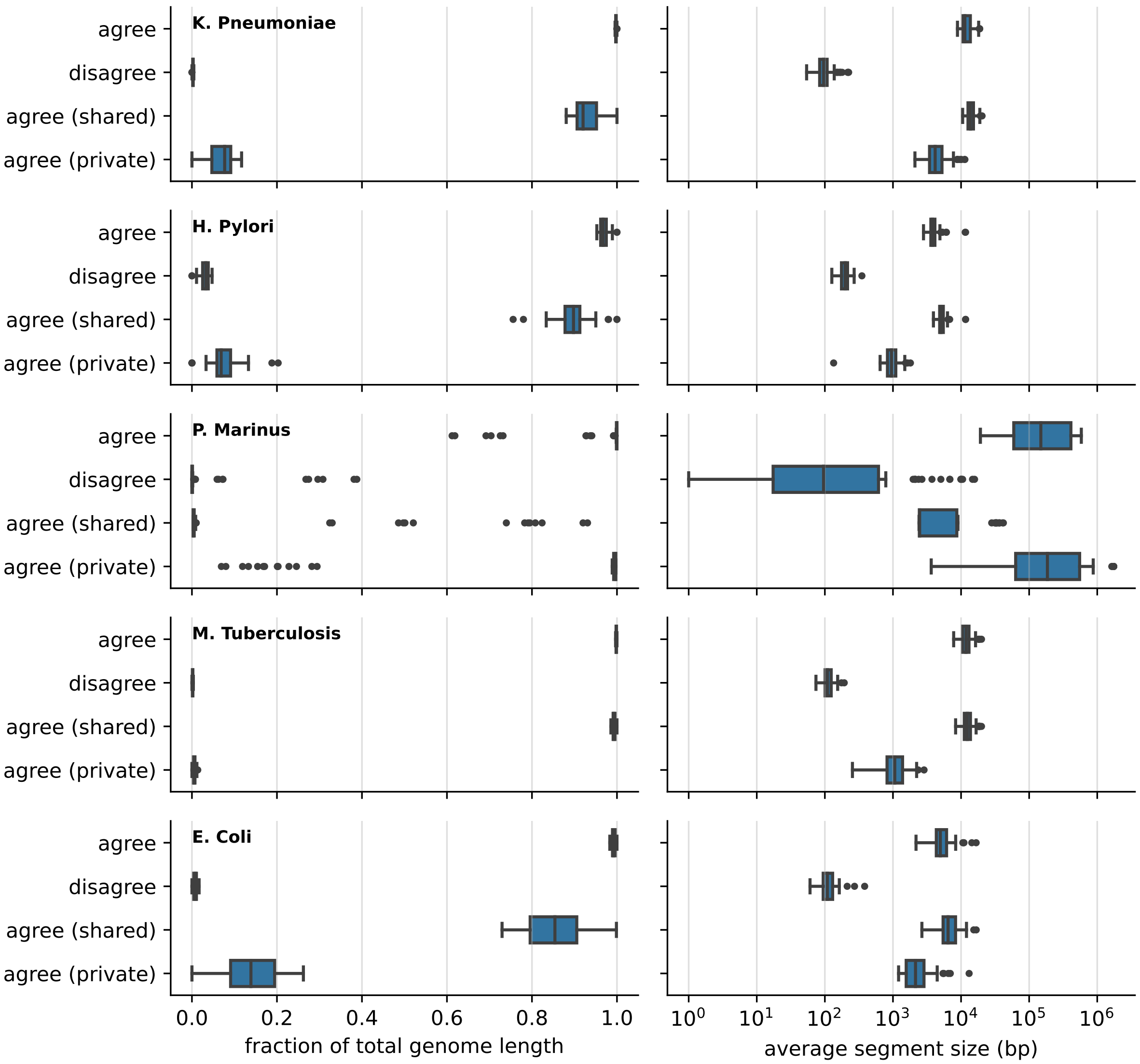


FIG. 5 **Marginalized graphs are compatible with pairwise graphs**. We compare marginalized graphs with pairwise built graphs for different random pairs of strains from 5 different species, as described in the text. Each graph defines a partition of a genome into segments, that can either be shared in the pair or private to the genome. We intersect these two partitions, and categorize segments based on whether the two partitions *agree* or *disagree* on their sharing. Agreement segments are further split depending on whether the segment is *shared* or *private* on both graphs. The figure displays the distribution of the average size of these segments (right column) and the average fraction of genome that they cover (left column) for the 50 different pairs considered (45 for PM). For all species with sequence divergence compatible with the limit of the alignment kernel used (KP, MT, EC), we find very good agreement between marginalized and pairwise graphs, with disagreement segment only covering a very small fraction of the genome and having size compatible with *PanGraph* threshold pancontig length (100 bp).
